# Epigenomic perturbation of novel *EGFR* enhancers reduces the proliferative and invasive capacity of glioblastoma and increases sensitivity to temozolomide

**DOI:** 10.1101/2023.02.22.529507

**Authors:** Craig A. Vincent, Itzel Nissen, Andreas Hörnblad, Silvia Remeseiro

## Abstract

Glioblastoma (GB) is the most aggressive of all primary brain tumours. Patients typically rely on radiotherapy with concurrent temozolomide (TMZ) treatment and face a median survival of ∼14 months. Alterations in the Epidermal Growth Factor Receptor gene (*EGFR*) are common in GB tumours, but therapies targeting EGFR have not shown significant clinical efficacy. Here, we investigated the influence of the *EGFR* regulatory genome on GB cells, and identified novel *EGFR* enhancers located in an intronic region nearby the GB-associated SNP rs723527. Epigenomic perturbation of this regulatory region using CRISPR-based methods decreases *EGFR* expression and reduces the proliferative and invasive capacity of glioblastoma cells, while increasing their sensitivity to TMZ. The enhancer-perturbed GB cells also undergo a metabolic reprogramming in favour of mitochondrial respiration and present increased apoptosis. Our findings demonstrate how epigenomic perturbation of *EGFR* enhancers can ameliorate the aggressiveness of glioblastoma cells and enhance the efficacy of TMZ treatment.

**SIGNIFICANCE:** Our study demonstrates how CRISPR/Cas9-based perturbation of enhancers can be used to modulate the expression of key cancer genes, which can help improve the effectiveness of existing cancer treatments and potentially the prognosis of difficult-to-treat cancers such as glioblastoma.

## INTRODUCTION

Glioblastoma (GB), also known as grade 4 astrocytoma, is a common and highly aggressive type of primary brain tumour, for which survival rates have not significantly improved in recent decades (median survival ∼14 months, 5-year survival <5%) (1,2). Due to its highly invasive nature, complete surgical resection is nearly impossible (3) and therefore recurrence is inevitable. Radiotherapy with concomitant chemotherapy, often preceded by tumour debulking surgery, forms the basis of GB standard of care. Temozolomide (TMZ) is a DNA alkylating agent commonly used in the treatment of glioblastoma as adjuvant to radiotherapy. Patients with methylated *MGMT* (O6-methylguanine-DNA methyltransferase) promoter respond with better outcome to TMZ treatment given the role of MGMT in DNA damage repair (4–6), which highlights the relevance of epigenomic cues in the patient outcome upon treatment.

In fact, GB remains a difficult-to-treat cancer due to the high degree of inter- and intra-tumour heterogeneity and the complexity of genetic, epigenetic and microenvironment events. One of the key challenges in treating glioblastoma is the ability of cancer cells to evade the effects of chemotherapy and radiation therapy. This is often due to the over-expression of certain genes, such as *EGFR*, which can promote the survival and proliferation of cancer cells. The Epidermal Growth Factor Receptor gene (*EGFR*) is one of the most frequently altered genes in glioblastoma. 57% of tumours display some form of alteration in *EGFR* (7) and among the classical subtype, *EGFR* is overexpressed in more than 95% (8). Moreover, high *EGFR* expression in gliomas correlates with reduced overall survival in patients (9). Constitutive activation of the EGFR signalling pathway can occur through overexpression of the receptor itself or its ligand, through amplification of the *EGFR* locus (which includes non-coding regions), or through coding mutations (e.g. EGFRvIII). All of which result in increased cell proliferation, invasive capacity, survival and angiogenic potential.

While the traditional focus of cancer research has been on the impact of coding mutations, Genome-Wide Association Studies (GWAS) have revealed that most genetic variants that predispose to cancer are located within non-coding genomic regions with potential to act as cis-regulatory elements (e.g. enhancers) (10). Enhancers are stretches of DNA that regulate transcription in a spatiotemporal manner, through their capacity to bind transcription factors (TFs) and protein complexes that control gene expression. In the linear genome, enhancers can be located vast distances from the gene promoter which they act upon, but they require close physical proximity in the 3D nuclear space to exert their regulatory function (11–13). Enhancer dysfunction due to genetic, topological or epigenetic mechanisms can contribute to human diseases, including cancer. However, accurate identification of enhancers and understanding their role in disease still remains a challenge (14).

In the context of glioblastoma, the mechanistic contribution of the non-coding regulatory genome to pathogenesis remains understudied. Here, we identify novel *EGFR* enhancer elements in the vicinity of the known GB-associated single nucleotide polymorphism (SNP) rs723527, and we functionally dissect their regulatory potential by introducing CRISPR-based (epi-)genomic perturbations. Targeting these *EGFR* enhancer regions in glioblastoma cells leads to decreased proliferation and migration rates, due in part to an increased rate of apoptosis, which could be triggered by an underlying metabolic reprogramming of these cells. Thus, targeting these novel *EGFR* enhancers diminishes the malignancy of glioblastoma cells by reducing their proliferative and invasive capacity, and sensitising them to treatment with TMZ.

Our findings highlight the association between *EGFR* expression and temozolomide efficacy, and demonstrate how CRISPR/Cas9-based targeting of enhancers can be used to modulate the expression of key cancer genes. Combining (epi-)genomic perturbation of enhancers with existing cancer treatments can improve their effectiveness and subsequently the prognosis of glioblastoma and other cancers difficult to treat.

## RESULTS

### Identification of novel EGFR enhancers in glioblastoma

We first identified a panel of 10 conserved elements (CE1-CE10) as potential candidates to regulate the expression of *EGFR* in glioblastoma. This identification was based on sequence conservation and *GeneHancer* prediction to interact with the *EGFR* promoter, together with our previous data on distribution of active chromatin marks and chromatin accessibility (Chakraborty *et al* bioRxiv doi.org/10.1101/2022.11.16.516797) (Fig. 1A). Two SNPs associated with increased GB risk are located in the *EGFR* locus: rs723527, within intron 1, and rs75061358, ∼150kb upstream of the *EGFR* transcription start site (TSS) (15). Of these, rs723527 is located within one of these conserved elements (CE5), in a highly accessible region and enriched in the active enhancer mark H3K27ac in a panel of patient-derived glioblastoma cell lines (Fig. 1A and Chakraborty et al bioRxiv doi.org/10.1101/2022.11.16.516797). In contrast, rs75061358 is not located within one of the CEs and does not display any features indicative of enhancer activity. We made similar observations regarding the distribution of chromatin marks around these SNPs in U251 glioblastoma cells, as measured by ChIP-qPCR (Fig. 1B). In particular subregions CE5C and CE6B, which are proximal to the SNP rs723527, displayed enrichment of the active enhancer mark H3K27ac and depletion of the repressive mark H3K27me3.

**Figure 1.**
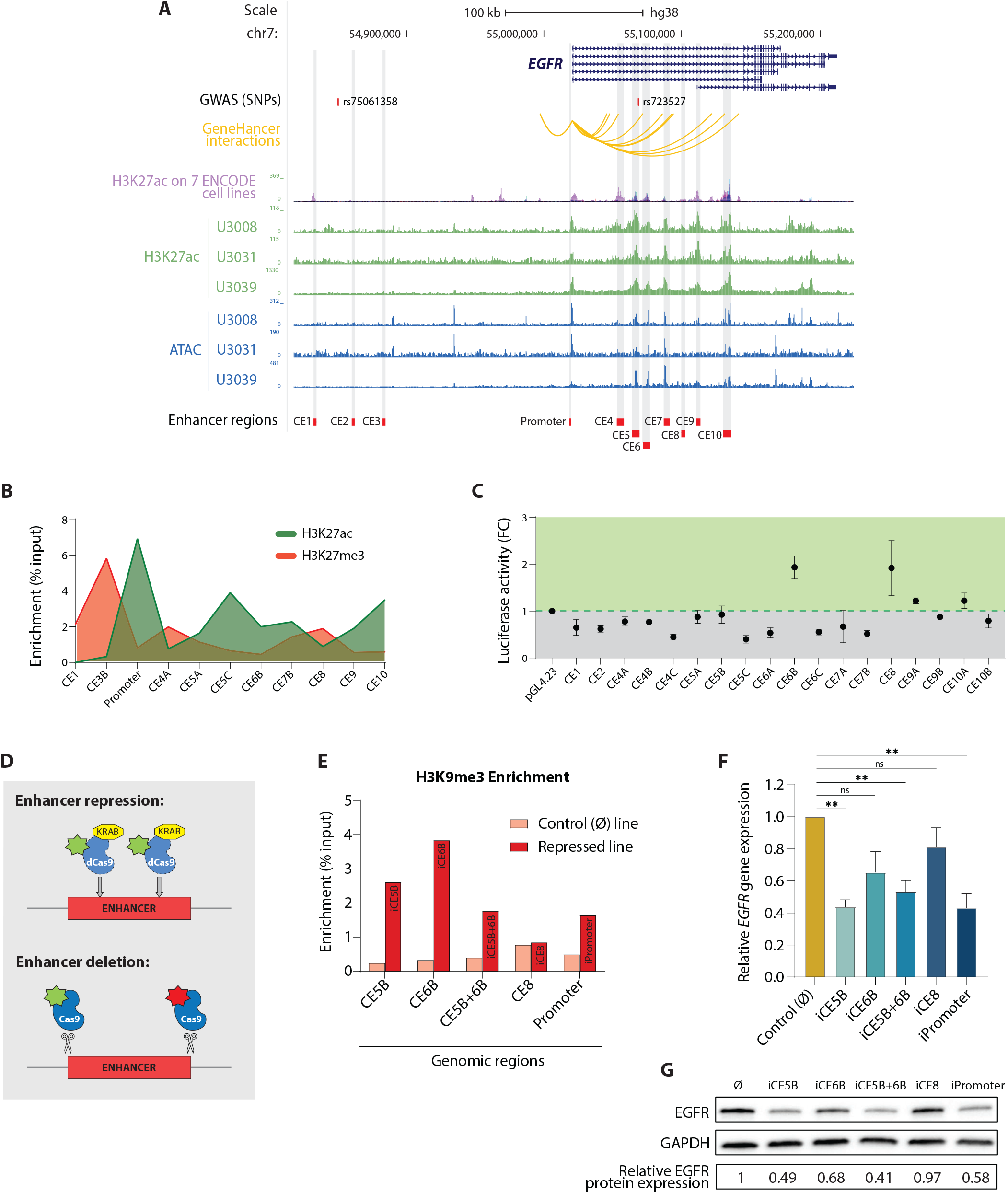
Identification of novel *EGFR* enhancers in glioblastoma located in the vicinity of the GB-associated SNP rs723527. **A**, Schematic representation of the *EGFR* gene locus displaying: GB-associated SNPs; *GeneHancer* predicted interactions between genomic regions and the *EGFR* promoter; H3K27ac enrichment in seven ENCODE cell lines; H3K27ac enrichment and chromatin accessibility by ATAC-seq across three representative patient-derived GB cell lines (our previous data Chakraborty et al bioRxiv doi.org/10.1101/2022.11.16.516797); and the conserved elements (CE) selected for characterisation highlighted in grey. Visualisation in the UCSC genome browser. **B**, Enrichment of H3K27ac and H3K27me3 in U251 glioblastoma cells around the CE regions as determined by ChIP-qPCR. **C**, Enhancer dual-luciferase reporter assay. Luciferase activity relative to the control reporter plasmid is expressed as a fold change. Data is presented as mean ± SEM (n=5). **D**, Schematic representation of the deletion and repression CRISPR-perturbation strategies. **E**, Enrichment of H3K9me3 upon expression of the transcriptional repressor KRAB in the CRISPRi-repressed cell lines as determined by ChIP-qPCR. **F**, *EGFR* gene expression relative to *HPRT* in each KRAB-repressed line (i.e. iCEx) as determined by RT-qPCR. Data is represented as mean ± SEM (n=4). Statistical significance was assessed by unpaired *t* test with Welch’s correction (** *P* < 0.01). **G**, EGFR protein expression determined by western blot and normalised to GAPDH protein levels.

To determine the regulatory potential of these 10 conserved elements (CEs), we employed luciferase reporter assays in U251 cells. For large CEs (>2kb), smaller regions were subcloned and tested (e.g. CE5 A, B and C regions). CE6B and CE8 retained the highest regulatory potential on enhancer reporter assays (Fig. 1C), where CE6B is located closest to the GB-associated SNP rs723527. In contrast, CEs located close to rs75061358 did not demonstrate enhancer activity in these reporter assays. These findings therefore highlight three putative enhancer elements: CE5, CE6 and CE8, which are located within intron 1 of *EGFR* and in close proximity to rs723527.

### CRISPR-perturbation of novel EGFR enhancers decreases EGFR gene expression and protein levels

To functionally demonstrate that the identified CEs act as *EGFR* enhancers in glioblastoma, we introduced targeted perturbations utilising both CRISPRi (dCas9-KRAB) and CRISPR/Cas9. We generated various U251 glioblastoma cell lines with either stable epigenomic repression of the CEs or carrying the deletions of interest (Fig. 1D). As expected, upon recruitment of the transcriptional repressor KRAB to the CEs, the established lines showed an enrichment of the repressive mark H3K9me3 in the corresponding region, in comparison to the empty vector control line (Fig. 1E, Supplementary Fig. S1A-F). dCas9-KRAB repression of the CEs (hereby iCE) correlated with significant downregulation of *EGFR* gene expression (Fig. 1F) and lower protein levels (Fig. 1G). For iCE5B, iCE5B+6B and the iPromoter region, *EGFR* gene expression levels were significantly reduced to 44%, 53% and 43% of the expression observed in the control line, respectively (Fig. 1F). Similarly, protein levels were reduced to 49%, 41% and 58% of the control line levels (Fig. 1G). Only in the case of iCE8, the level of repression indicated by enrichment of H3K9me3 was not sufficient to considerably diminish the EGFR protein levels. Furthermore, the cell lines carrying genomic deletions (Supplementary Fig. S2A) also present a significant downregulation of *EGFR* gene expression accompanied by reduced protein levels (Supplementary Fig. S2B, C). In the Δ CE5B+6B, Δ CE6B, ΔCE8 and ΔPromoter lines, *EGFR* expression is reduced to 29%, 48%, 66% and 70% of the control line levels, respectively (Supplementary Fig. S2B). Therefore, CRISPR-based perturbation of the CEs with regulatory potential demonstrates, in a functional manner, that they act as *EGFR* enhancers in the context of glioblastoma.

### Repressing the EGFR enhancers reduces the proliferative and invasive capacity of glioblastoma cells

Having determined the impact of enhancer perturbation on *EGFR* expression, we then evaluated the proliferative and invasive capacities of the enhancer-perturbed glioblastoma lines. Firstly, we assessed cell proliferation by live-cell imaging using the IncuCyte S3 live-cell analysis instrument and automated cell counting software. Cell lines with independent repression of CE5B and CE6B displayed a modest reduction in their proliferative capacity in comparison to the control (Fig. 2A). However, CRISPRi of the large region comprising CE5B+6B, which includes the SNP rs723527, significantly reduced the cell proliferation of glioblastoma cells to almost the same extent as the *EGFR* promoter-repressed cell line (Fig. 2A, B). CRISPR/Cas9-mediated deletion of the *EGFR* enhancers demonstrated slight inhibition of proliferation, though statistically insignificant, in all cell lines carrying the enhancer deletions (Supplementary Fig. S2D). The proliferative defect observed in the enhancer-repressed cell lines is much stronger than that of the enhancer-deletion lines, likely due to the spreading of the repressive marks over a larger region. Together with the added advantage that CRISPRi with dCas9-KRAB does not involve direct modification of the DNA sequence but solely epigenomic editing, we focused our further investigation on the *EGFR* enhancer-repressed glioblastoma cell lines.

**Figure 2.**
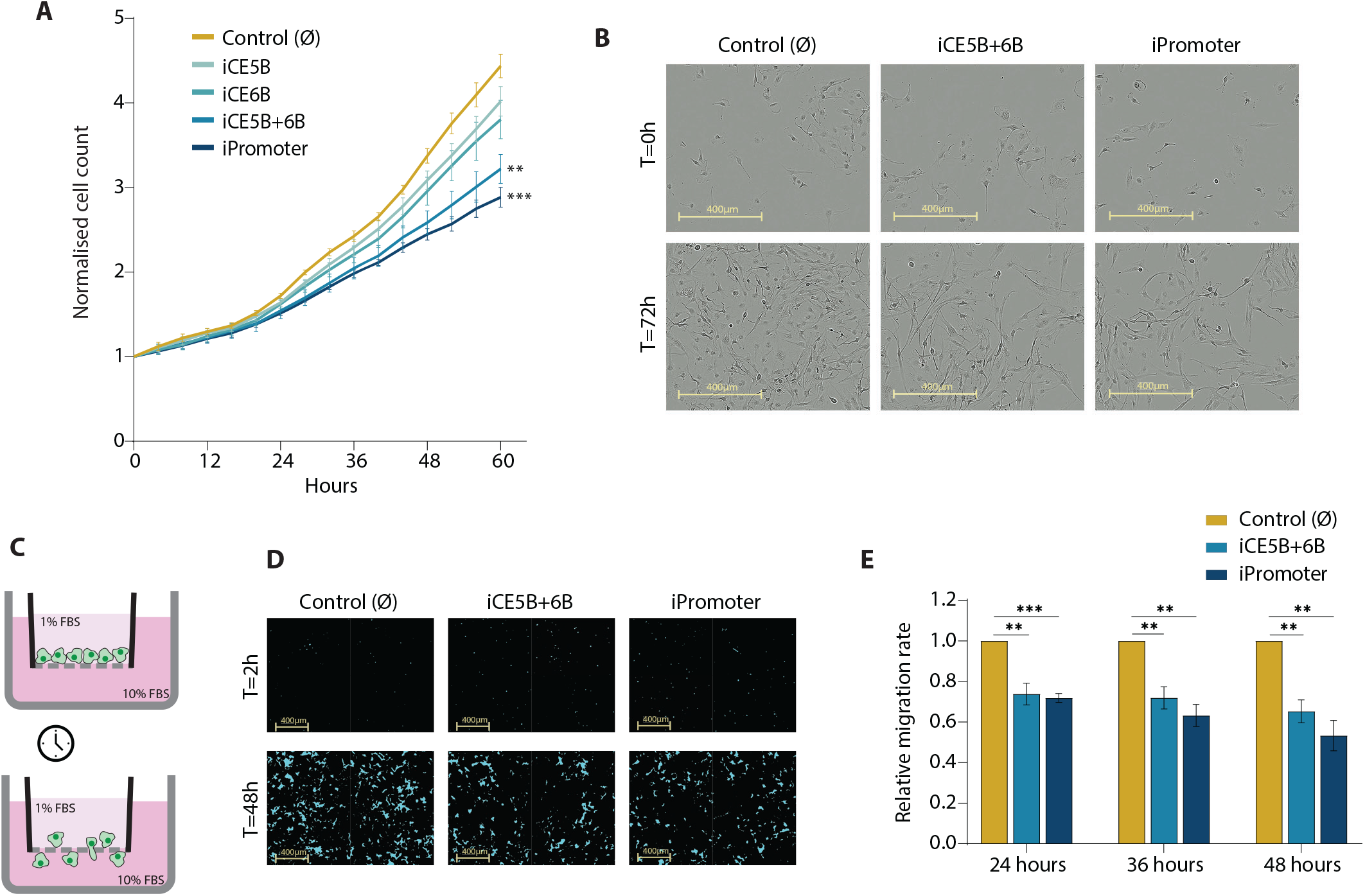
CRISPRi of novel *EGFR* enhancers reduces the proliferation and migration of glioblastoma cells. **A**, Proliferation rates of the *EGFR* enhancer-repressed lines determined by live-cell imaging. Images were acquired every 4 hours and proliferation was determined by automatic cell count. Data is normalised to t=0 and presented as mean ± SEM (n=4). Statistical significance was assessed by unpaired *t* test with Welch’s correction (** *P* < 0.01, *** *P* < 0.001). **B**, Representative images of control cells alongside iCE5B+6B and iPromoter cell lines at t=0h and t=72h. **C**, Schematic representation of the chemotactic migration assay. **D**, Representative images from the chemotactic assays taken at t=2h and t=48h. Masked area (blue) covers cells that migrated through the pores of the culture plate towards the chemoattractant. **E**, Relative migration rates of iCE5B+6B and iPromoter cell lines represented as total masked area of migrated cells at t=24h, t=36h and t=48h, normalised to the initial seeding density. Data is presented as mean ± SEM (n=3). P values were determined by unpaired *t* test (** *P* < 0.01, *** *P* < 0.001).

Next, we determined the invasive capacity of the *EGFR* enhancer-repressed lines by measuring their migration rate towards a chemical stimulus in chemotaxis assays. In these trans-well chemotactic assays, cells migrate through cell-permeable pores attracted by higher concentration of nutrients (i.e. from 1% to 10% FBS) and are monitored in real-time (Fig. 2C). As a negative control, a no-chemoattractant condition was established (i.e. 1% to 1% FBS) (Supplementary Fig. S3A-C). We observed that the migrative capacity of the iCE5B+6B cell line was significantly compromised (Fig. 2D-E), and similar to that observed upon repression of the *EGFR* promoter. Altogether, these findings show that repressing the *EGFR* enhancer region CE5B+6B, which encompasses the GB-associated SNP rs723527, leads to significantly decreased proliferation and migration of glioblastoma cells.

### Reduced malignancy of the EGFR-enhancer repressed GB cells can be linked to increased apoptosis and mitochondrial respiration

We further characterised the *EGFR-*enhancer repressed lines by firstly measuring their relative apoptosis rates over time by live-cell imaging of cell cultures in the presence of annexin V red dye. The apoptotic cells (i.e., annexin V positive area) in the iCE5B+6B repressed GB line increased at a significantly faster rate compared to the control cell line (Fig. 3A, D). The rate of apoptosis within the individually repressed CE5B and CE6B cell lines does not significantly differ from the unmodified control cells, and interestingly, nor does the promoter-repressed cell line. This effect is also observed when we account for differences in proliferation rate by normalising the annexin V-positive area to the total cell population area. We observed that after 48 and 72 hours in culture, the percentage of annexin V-positive area in the iCE5B+6B cell line is significantly higher than that of the control line (Fig. 3B, C). This suggests that targeting the CE5B+6B enhancer region specifically causes an apoptotic response which cannot be triggered by repressing the promoter of *EGFR*.

**Figure 3.**
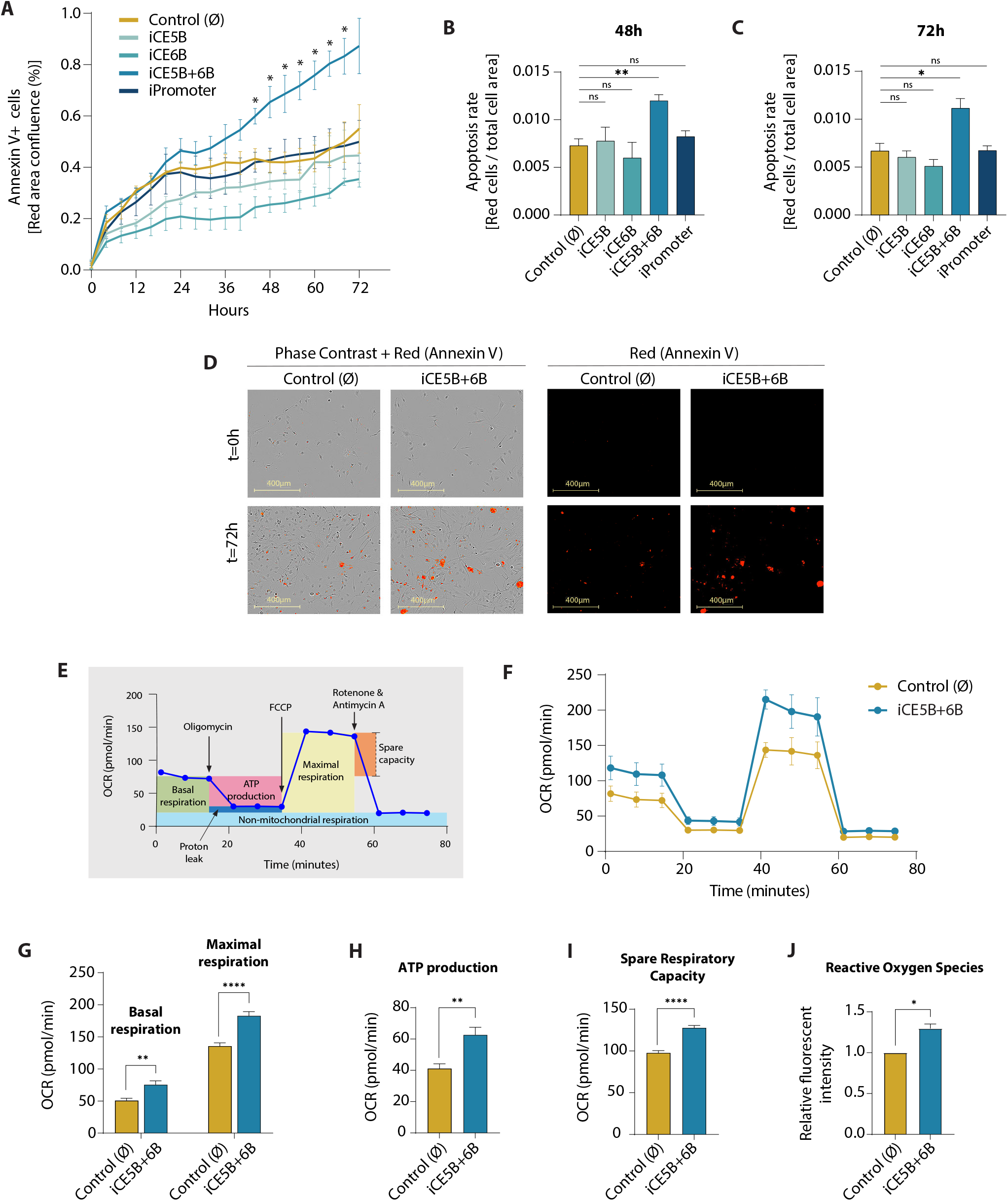
Epigenomic perturbation of the *EGFR* enhancer CE5B+6B triggers apoptosis and favours mitochondrial respiration. **A**, Apoptosis levels in the *EGFR* enhancer-repressed lines as determined by annexin V red fluorescence area (% confluence) measured at 4-hour intervals. Data is presented as mean ± SEM (n=3). P values were determined by unpaired *t* test with Welch’s correction (* *P* < 0.05). **B-C**, Apoptosis rate represented as proportion of the area occupied by annexin V red apoptotic cells *vs* total cells at t=48h (**B**) and t=72h (**C**). Data is presented as mean ± SEM (n=3). Statistical significance was determined by unpaired *t* test (* *P* < 0.05, ** *P* < 0.01). **D**, Representative phase-contrast images of control cells and iCE5B+6B cells alongside annexin V-positive cells (red) to identify apoptotic cells at t=0h and t=72h. **E**, Schematic representation of the Agilent Seahorse XF Cell Mito Stress Test. **F**, Oxygen Consumption Rate (OCR) of control cells and iCE5B+6B enhancer-repressed cells in response to the assay compounds. Data is plotted as mean ± SEM (n=3). **G-I**, Basal and maximal respiration (**G**), ATP production (**H**) and spare respiratory capacity (**I**) of control cells and iCE5B+6B enhancer-repressed cells as determined by Cell Mito Stress Test. *P* values were determined by unpaired *t* test (** *P* < 0.01, **** *P* < 0.0001). **J**, Levels of reactive oxygen species (ROS) in control and iCE5B+6B cells represented as integrated red fluorescent intensity per cell count. Data is presented as mean ± SEM (n=3). Statistical significance assessed by unpaired *t* test with Welch’s correction (* *P* < 0.05).

Cancer cell metabolism is a key factor contributing to the cells’ ability to evade apoptosis. In order to examine whether this increased rate of apoptosis observed in the iCE5B+6B GB line was linked to changes in cellular metabolism, we performed a Seahorse Cell Mito Stress Test (Fig. 3E, F) to measure the relative oxygen consumption rates (OCR) of the cell lines as an assessment of mitochondrial function. We found that the iCE5B+6B-repressed line presents a significantly higher basal and maximal OCR compared to the control line (Fig. 3G). This would suggest that these *EGFR-*enhancer repressed cells are favouring mitochondrial respiration over glycolysis. Based on the same assay, we can also extract that the ATP production and spare respiratory capacity (SRC) of the enhancer-repressed line increased significantly over the control (Fig. 3H, I). Moreover, the increased mitochondrial respiratory parameters in the iCE5B+6B cell line are accompanied by significantly increased production of ROS (Reactive Oxygen Species) (Fig. 3J). These findings indicate that epigenomic perturbation of the CE5B+6B enhancer region causes increased mitochondrial respiration, resulting in an increased production of ROS, which would contribute to the apoptotic response observed.

### Epigenomic perturbation of the EGFR enhancers sensitises glioblastoma cells to TMZ treatment

Since temozolomide (TMZ) is the first-choice chemotherapeutic agent to treat GB clinically, we wanted to address how the *EGFR-*enhancer repressed lines respond to treatment with the drug. Not only the combined iCE5B+6B, but also the individual iCE5B, iCE6B and the *EGFR* iPromoter lines, showed a significantly slower proliferation rate upon TMZ treatment than the DMSO-treated controls (Fig. 4B-E and F). On the contrary, the empty vector control line is not significantly affected by TMZ treatment at the used concentration (Fig. 4A, F). Therefore, epigenomic repression of *EGFR* regulatory elements (i.e., novel enhancers and promoter), and subsequent downregulation of *EGFR* gene expression, sensitises glioblastoma cells to TMZ treatment. Our results show that combining epigenomic perturbation of enhancers or gene promoters with existing cancer drugs could improve the effectiveness of current treatments and subsequently the prognosis of patients.

**Figure 4.**
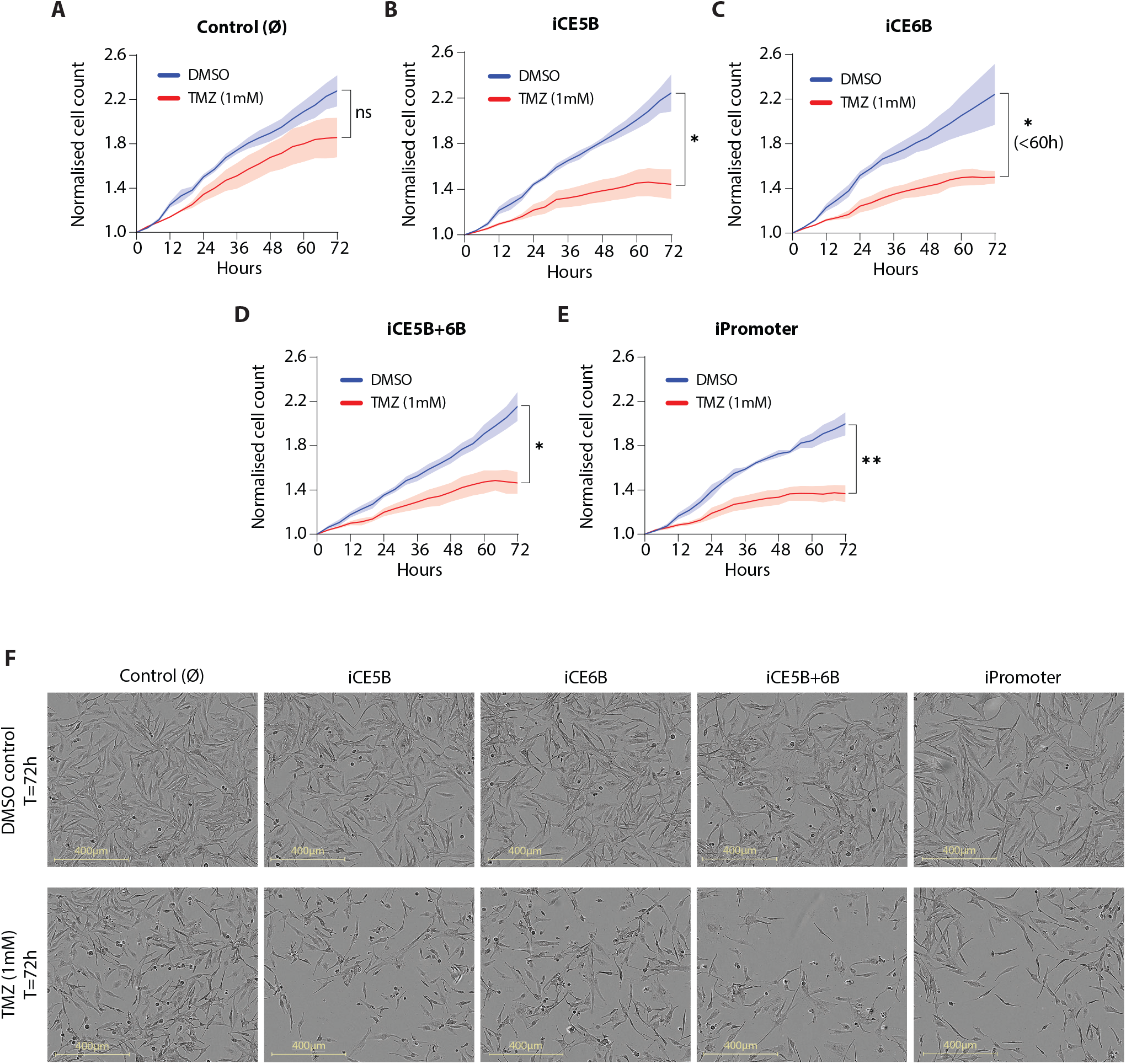
Epigenomic repression of the novel *EGFR* enhancers sensitises glioblastoma cells to temozolomide (TMZ) treatment. **A-E**, Proliferation rates of the *EGFR* enhancer-repressed lines determined by live-cell imaging upon treatment with 1mM TMZ in comparison with the DMSO-treated control. Images were acquired every 4 hours and proliferation was determined by automatic cell count. Data is normalised to t=0h and represented as mean ± SEM (n=3). *P* values were determined by unpaired *t* test (* *P* < 0.05, ** *P* < 0.01.) **F**, Representative images of iCE5B+6B, iPromoter and control cells upon TMZ treatment in comparison to DMSO-treated controls at t=72h.

## DISCUSSION

This study identified novel enhancers that drive the expression of *EGFR* in glioblastoma cells. CRISPR-mediated (epi-)genomic perturbation (i.e., repression, deletion) of these enhancer regions has a direct effect on the survivability and invasiveness of glioblastoma cells. By specifically repressing the CE5B+6B enhancer region that encompasses the known GB-associated SNP rs723527, we can lower *EGFR* expression levels and modulate the aggressiveness of U251 glioblastoma cells, which become less proliferative and invasive.

One underlying component of this is an apparent shift in the cellular metabolism upon enhancer perturbation and subsequent *EGFR* downregulation. The *EGFR* iCE5B+6B cells increase their basal and maximal mitochondrial respiratory activity, indicating a shift from the typical preference for glycolysis that is a common hallmark of cancerous cells (16–18). Higher mitochondrial respiration rates result in greater production of reactive oxygen species (ROS), which in turn can inhibit cell growth, damage cellular components and induce cell death (19). Deregulation of ROS production and ROS limitation pathways are common features of cancer cells (20). The metabolic rewiring in favour of mitochondrial respiration that we observe in the *EGFR* iCE5B+6B cells is accompanied by an increased accumulation of ROS and, subsequently, an increase in apoptotic events. This ultimately contributes to a reduction of cell proliferation upon repression of the *EGFR* enhancers in glioblastoma.

Migration of cancer cells in response to chemical stimuli is an important mechanism in the tumour dissemination process, both locally and during metastatic progression (21). The tumour-associated microglia and macrophages (TAMs) present in the GB tumour microenvironment release growth factors and cytokines, including EGF (Epidermal Growth Factor) and CSF-1, which can promote tumour proliferation, survival and invasion (22,23). Our *EGFR* enhancer-repressed glioblastoma cells also present a reduced response to chemo-attractive stimuli and express less *EGFR* than the parental unmodified cells. One could therefore speculate that *in vivo* they might be less responsive to EGF being secreted by macrophages in the tumour microenvironment and could therefore be less invasive.

Repressing the CE5B+6B *EGFR* enhancer reduces the proliferative and invasive capacity of GB cells, therefore ameliorating the malignant phenotype of glioblastoma cells, while additionally sensitising the cells to temozolomide: the current chemotherapeutic of choice in the clinic. The nature of the relationship between *EGFR* amplification levels and the response to TMZ treatment remains inconclusive and under debate (24). In our study, upon enhancer repression, lower *EGFR* levels correlate with an improved response to TMZ. Our findings point to an increased effect of temozolomide in combination with *EGFR* enhancer perturbation that may provide an effective combination therapy.

Taken together, our data highlights the functional importance of the *EGFR* regulatory genome in glioblastoma and it demonstrates the potential of enhancer modulation as a therapeutic strategy. In the future, the combination of epigenomic perturbation of enhancers and current anti-cancer drugs can improve their effectiveness and subsequently the prognosis of difficult-to-treat cancers, such as glioblastoma.

## METHODS AND MATERIALS

### Cell Culture

*U251 glioblastoma cells* (Sigma-Aldrich, #09063001, authenticated by short tandem repeat (STR)-PCR profiling) were grown in EMEM (EBSS) supplemented with 2mM Glutamine, 1% NEAA (Non-Essential Amino Acids), 1mM Sodium Pyruvate, 10% FBS (Fetal Bovine Serum) and 1% penicillin/streptomycin (all from Gibco). HEK293T cells were grown in DMEM/F-12 GlutaMAX™-Supplemented media containing 10% FBS (Fetal Bovine Serum) and 1% penicillin/streptomycin (all from Gibco). All cell lines were grown in a cell incubator at 37°C in a humidified atmosphere (95% humidity) with 5% CO2.

### Luciferase Dual-Reporter Assay

Luciferase assay was performed using the Promega Dual-Luciferase® Reporter Assay System following manufacturer’s instructions. Conserved Elements (CE) were PCR-amplified from GB genomic DNA using GoTaq® G2 DNA Polymerase (Promega, #M7845) (primer sequences listed in Supplementary Table S1) and cloned into pGL4.23[luc2/minP] vector (#E8411, Promega) using Acc65I and BglII restriction enzymes (Thermo Fisher). pGL4.23+enhancer constructs were transfected into the U251 cells using Lipofectamine™ 2000 Transfection Reagent (Thermo Fisher) together with the pRL-SV40 vector (#E2231, Promega) for signal normalisation. pGL4.13[luc2/SV40] vector (#E6681, Promega) served as a positive control. Luminescence readings were taken using the Biotek Synergy HT microplate reader. Data was represented as fold change (FC) over empty pGL4.23 readings.

### ChIP (Chromatin ImmunoPrecipitation)-qPCR

ChIP-qPCR was performed in stable glioblastoma lines established upon epigenomic perturbation of the *EGFR* enhancers, including the empty vector control lines, and in the parental U251 GB cell line. Briefly, cells were fixed on the plate by adding formaldehyde directly to the medium (final concentration 1% formaldehyde) for 15 minutes at room temperature while rotating. The crosslinking reaction was quenched by adding Glycine (final concentration 125mM Glycine) for 5 min, and fixed cells were scraped off and harvested in 1X cold PBS containing protease inhibitors. Cells were then resuspended in lysis buffer (3-6×10^6^ cells/ml) and sonicated in a Covaris E220 instrument (shearing time 12min, PIP 140, duty factor 5, 200 cycles per burst). Chromatin immunoprecipitation was performed with antibodies against H3K27ac (Abcam Cat# ab4729), H3K27me3 (Abcam Cat# ab192985) and H3K9me3 (Abcam Cat# ab8898) and using Dynabeads™ M-280 Sheep Anti-Rabbit IgG (Invitrogen Cat# 11203D). Chromatin Immunoprecipated DNA was amplified by qPCR using a CFX Connect Real-Time PCR Detection System (Bio-Rad) and primers specific for the genomic regions of interest (Supplementary Table S2). Positive and negative regions were measured in parallel for control purposes and enrichment is calculated over the input.

## Generation of Stable Cell Lines

### Cloning

The UCSC genome browser tool ‘CRISPR target identifier’ was used to select CRISPR gRNAs. For CRISPRi, gRNAs targeting central regions of the CEs were cloned into the *pLV hU6-sgRNA hUbC-dCas9-KRAB-T2a-GFP* plasmid (Addgene, #71237) using the BsmBI restriction sites (gRNA sequences listed in Supplementary Table S3). To generate genomic deletions, we modified this plasmid and cloned gRNAs targeting the flanks of the CEs. First, we replaced the dCas9-KRAB with an active Cas9 coding sequence, and further replaced GFP by mCherry, thus generating two new constructs hereby named *pLV hU6-sgRNA hUbC-Cas9-T2a-GFP* and *pLV hU6-sgRNA hUbC-Cas9-T2a-mCherry*. Deletion gRNAs were then cloned into these vectors. The GFP and mCherry expression enabled subsequent FACS sorting of positively transduced cells.

### Lentivirus transduction

Lentiviral particles were produced and collected upon transfection of HEK293T cells with the lentiviral Cas9 or dCas9 plasmids expressing the gRNAs, along with the pSPAX and pMD2.G lentiviral packaging plasmids and using Lipofectamine™ 2000 (Thermo Fisher). Between 24-48 hours post-transfection, the viral supernatant was filtered, supplemented with 20mM HEPES and polybrene (10μg/ml), and used for transduction of U251 cells in three rounds.

### FACS Sorting

To establish stable lines, transduced cells were sorted by Fluorescence-Activated Cell Sorting (FACS) using the BD FACSAria™ III Cell Sorter instrument and the BD FACSDiva software. For CRISPRi experiments, GFP positive cells were collected and, in the case of CRISPR/Cas9-mediated genomic deletions double positive GFP+mCherry+ cells were sorted and further expanded.

### Validation of cell lines

Repression by CRISPRi was validated by measuring the enrichment of H3K9me3 by ChIP-qPCR (see methods section above). Genomic deletions were confirmed by genotyping PCR using primers designed to flank the gRNA target sequences. Genomic DNA (gDNA) was extracted using Qiagen DNeasy Blood & Tissue Kit (ID: 69504) and genotyping PCRs were performed using GoTaq® G2 DNA Polymerase (Promega, #M7845) (genotyping primers listed in Supplementary Table S4).

### RT-qPCR

Total RNA was extracted from cells using the RNeasy Plus Mini Kit (ID: 74134, Qiagen). cDNA was synthesised using RevertAid H Minus Reverse Transcriptase (#EP0451, Thermo Fisher) and random hexamers (#SO142, Thermo Fisher), following the manufacturer’s instructions. Quantitative-PCR analysis was performed with CFX Connect Real-Time PCR Detection System (Bio-Rad) using SYBR green master mix - PowerUp (Thermo Fisher). *EGFR* gene expression was measured alongside the housekeeping gene *HPRT* for normalization (qPCR primers listed in Supplementary Table S5). Relative expression levels were determined using the Δ ΔCt method.

### Western blot

Whole cell protein extracts were prepared using lysis buffer containing 20% SDS and 1M Tris-HCl pH 6.8. Protein concentration was measured using the Pierce™ BCA Protein Assay Kit (Thermo Scientific) and absorbance at 560nm was determined using the Biosan HiPo MPP-96 microplate photometer. Protein samples were loaded into precast gels, run in the Mini-PROTEAN Tetra Cell and blotted using the Trans-Blot® Turbo™ Transfer System (all Bio-Rad) according to standard protocols. Primary antibodies against EGFR (1:1000, rabbit, Cell Signaling Cat# 4267) and GAPDH (1:1000, rabbit, Cell Signaling Cat# 2118) were diluted in 5% bovine serum albumin (BSA) in Tris-Buffered Saline 0.1% Tween® 20 Detergent. An HRP-conjugated goat anti-rabbit secondary antibody (1:10000, Jackson ImmunoResearch Labs Cat# s111-035-003) was used for detection together with Bio-Rad Clarity Western ECL Substrate. ChemiDoc™ MP Imaging System with Image Lab™ Software (Bio-Rad) was used for signal detection and quantification.

### Live-Cell Imaging

All live-cell imaging experiments were performed using the IncuCyte S3 Live-Cell Analysis instrument (Sartorius) and the image analysis was performed using the Incucyte Base Analysis Software.

### Proliferation assays

Cell proliferation was determined by live-cell imaging taking phase-contrast images every 4 hours during a period of 72 hours. Automated cell segmentation and counting was performed with the adherent Cell-by-Cell analysis software module. Data was normalised to the t=0h count and presented as ratios.

### Chemotaxis assays

Chemotactic migration was determined by imaging cells in the Incucyte® Clearview 96-Well Chemotaxis Plate (#4582), and analysed using the Chemotaxis Analysis Software Module. Cells were seeded in 1%FBS media in the trans-well insert and 10%FBS media was used as chemoattractant in the reservoir wells. A no-chemoattractant negative control was set up using 1% FBS in both the insert and reservoir wells.

### Annexin V Apoptosis assays

Incucyte® Annexin V Red Dye (Sartorius #4641) was added to the cell culture medium at a final dilution of 1:200 (as per product guidelines). Both phase-contrast and red fluorescence (Excitation: 567–607nM, Emission 622–704nM) images were taken every 4 hours during a period of 72 hours. A red area confluence mask was applied to the cells to measure the apoptotic cell area using the Incucyte Base Analysis Software. Data was expressed as red area confluence (%) and normalized to total cell count (red area/total phase area).

### Reactive Oxygen Species

5μM CellROX™ Deep Red Reagent (Invitrogen #C10422) was added to cells in culture. After 30 minutes of incubation time at 37°C, the reagent was washed out twice with PBS and the cells were immediately imaged. Both phase-contrast and red fluorescence (Excitation: 567–607nM, Emission 622–704nM) images were taken. A mask was applied to the red fluorescent signal to measure integrated intensity (normalised to phase-contrast cell count).

### Temozolomide treatment

Cells were treated with 1mM TMZ (Temozolomide, Sigma-Aldrich T2577) dissolved in DMSO (Dimethyl Sulfoxide, Calbiochem - CAS 67-68-5) and cell proliferation was assessed in comparison to DMSO-treated control cells as above (see *Proliferation assays*).

### Measurement of Mitochondrial Function

Mitochondrial function was determined using the Seahorse XFe96 Analyzer (Agilent), which measures mitochondrial oxygen flux and extracellular acidification rate for live cells in real time. The Cell Mitochondrial Stress Test was performed following manufacturer’s instructions and oxygen consumption rates (OCR) were determined. Seahorse 96 well-plates were coated with Poly-D-lysine (50μg/ml) and 20,000 cells were seeded per well. The test was performed as per standard protocol in XF assay medium (Dulbecco’s Modified Eagle Medium (DMEM) + 5mM glucose + 2mM glutamine + 1Mm pyruvate, pH7.4). 20μM of oligomycin, 10μM of FCCP and 5μM rotenone + 5μM Antimycin A were added to selectively inhibit different steps of mitochondrial respiration and thus initiate the relevant phases of the test. ATP production was calculated as (basal respiration – proton leak). Spare Respiratory Capacity (SRC) was determined as (maximal respiration – basal respiration).

### Statistical analysis

All statistical analysis was performed using GraphPad Prism 9 software. Statistical tests, number of replicates and significance are indicated in figure legends and in the corresponding figure panels.

## AUTHORS’ CONTRIBUTIONS

CA-V designed, performed, analysed and interpreted most of the experiments and wrote the original manuscript draft. IN performed ChIP-qPCR experiments. AH contributed to data interpretation and manuscript editing. SR designed and supervised the study, secured funding, analysed and interpreted the data, and wrote and edited the final manuscript with input from the other authors.

## AKNOWLEDGEMENTS

We acknowledge the Biochemical Imaging Center at Umeå University and the National Microscopy Infrastructure, NMI (VR-RFI 2019-00217) for providing assistance in microscopy, and the Umeå University Flow Cytometry platform for assistance with cell sorting. Seahorse analyses were performed at the Seahorse Platform, Umeå University. We also thank Helena Edlund for providing the pSPAX and pMD2.G plasmids. Research in SR’s laboratory is supported by the Swedish Research Council (2019-01960), the Swedish Cancer Foundation (21 1720), Knut och Alice Wallenbergs Stiftelse (WCMM), the Kempe Foundation (SMK-1964.2) and the Cancer Research Foundation Norrland (AMP 19-977, LP 21-2290, AMP 22-1091).

---

**Supplementary Figure 1.**
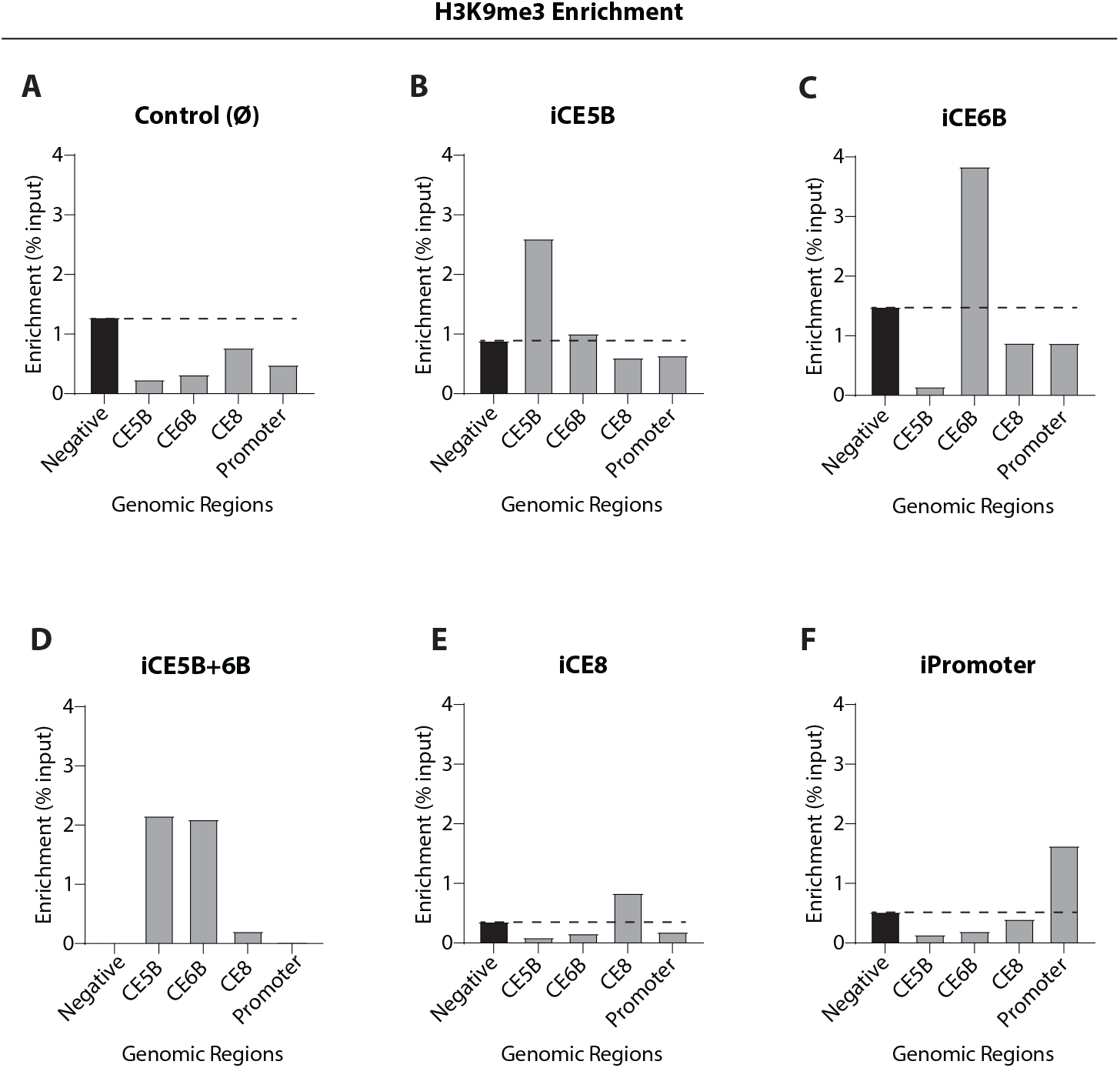
Recruitment of dCas9-KRAB repressor complex leads to enrichment of H3K9me3 at specific targeted sites. **A-F**, Bar charts depicting the enrichment of H3K9me3, as determined by ChIP-qPCR, at each genomic region (i.e. CE5B, CE6B, CE8, Promoter) and in each of the *EGFR* enhancer-repressed lines: iCE5B (**B**), iCE6B (**C**), iCE5B+6B (**D**) and iCE8 (**E**), alongside the iPromoter (**F**) and control line (**A**).

**Supplementary Figure 2.**
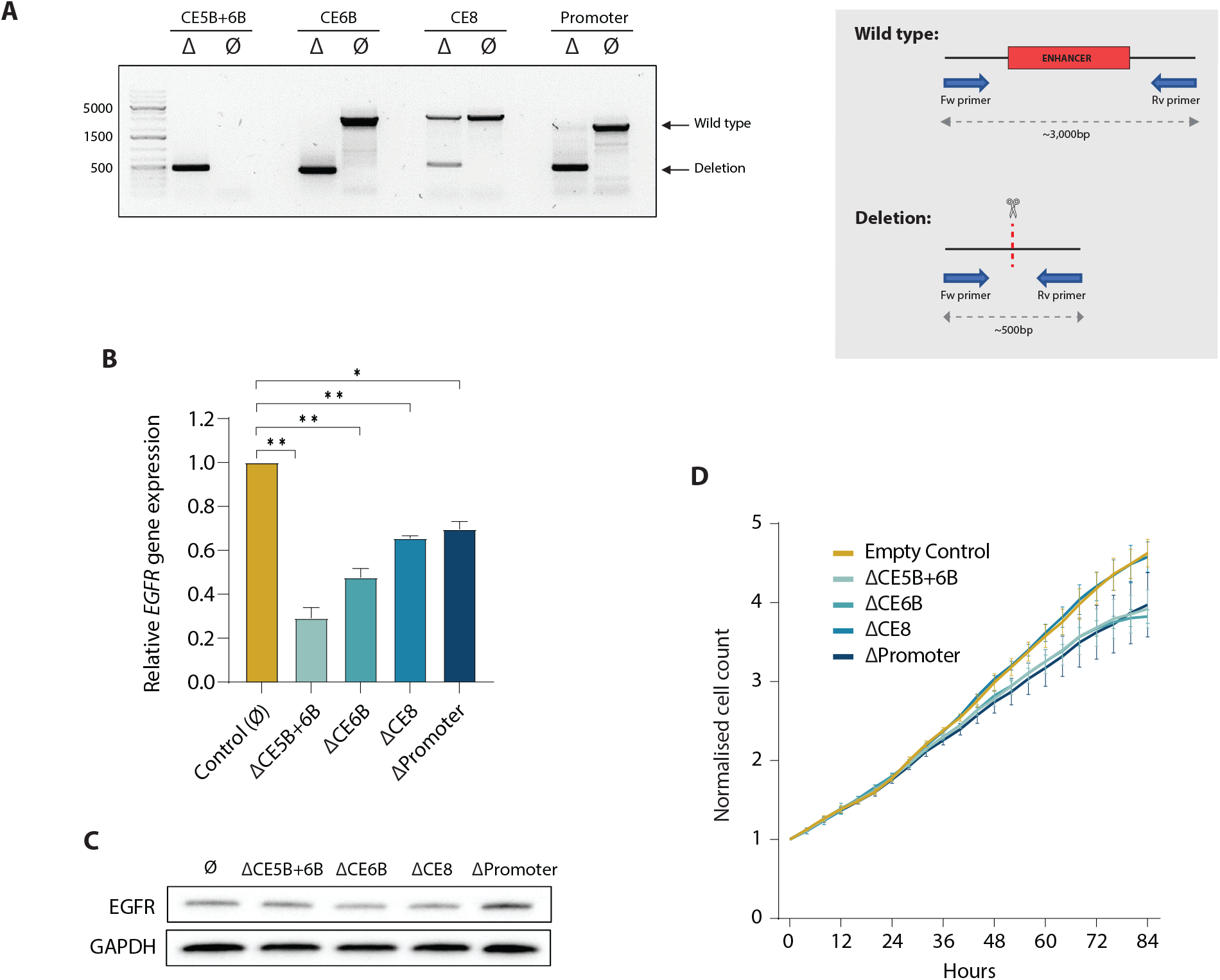
CRISPR/Cas9-mediated deletion of *EGFR* enhancers downregulates *EGFR* gene expression and affects cell proliferation rates. **A**, Genotyping PCR of the *EGFR* enhancer-deleted cell lines (ΔCE5B, ΔCE6B, ΔCE5B+6B, ΔCE8) alongside the ΔPromoter and empty vector control lines (left), and schematic outline of the PCR genotyping strategy (right). Note that the wild-type CE5B+6B allele is too large to be amplified under these conditions. **B**, *EGFR* gene expression levels relative to *HPRT* in *EGFR* enhancer-deleted cell lines as determined by RT-qPCR assays. Data is represented as mean ± SEM (n=3). Statistical significance as assessed by unpaired *t* test with Welch’s correction (* *P* < 0.05, ** *P* < 0.01). **C**, EGFR protein expression determined by western blot and normalised to GAPDH protein levels. **D**, Proliferation rates of cell lines carrying *EGFR* enhancer deletions or promoter deletions as determined by live-cell imaging, and in comparison to the empty vector control line. Images were acquired every 4 hours and proliferation was determined by automatic cell count. Data is normalised to t= 0h and plotted as mean ± SEM (n=3).

**Supplementary Figure 3.**
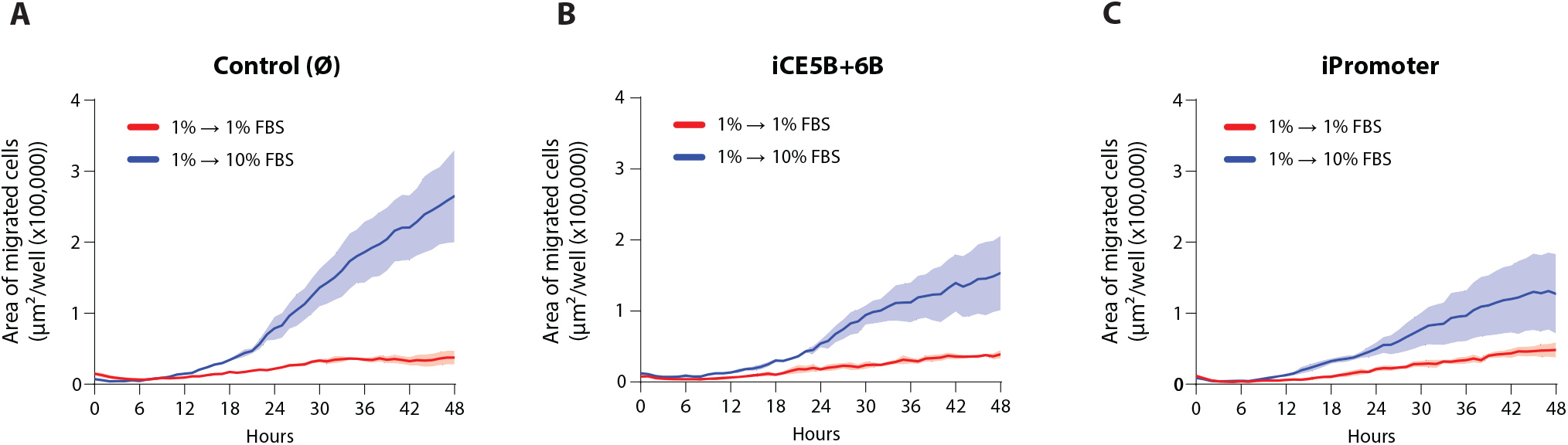
CRISPRi of the *EGFR* enhancer CE5B+6B and promoter compromises the migration of glioblastoma cells. **A-C**, Line plots comparing the rate at which the respective cell lines migrate either from media containing 1%FBS to 1%FBS (no-chemoattractant negative control) or from 1%FBS to 10%FBS (chemoattractant condition). Migration was assessed by live-cell imaging taking images every hour and migration rate was determined by automatic quantification of the area of migrated cells. Data is represented as mean ± SEM (n=3).

**Supplementary Table S1.**
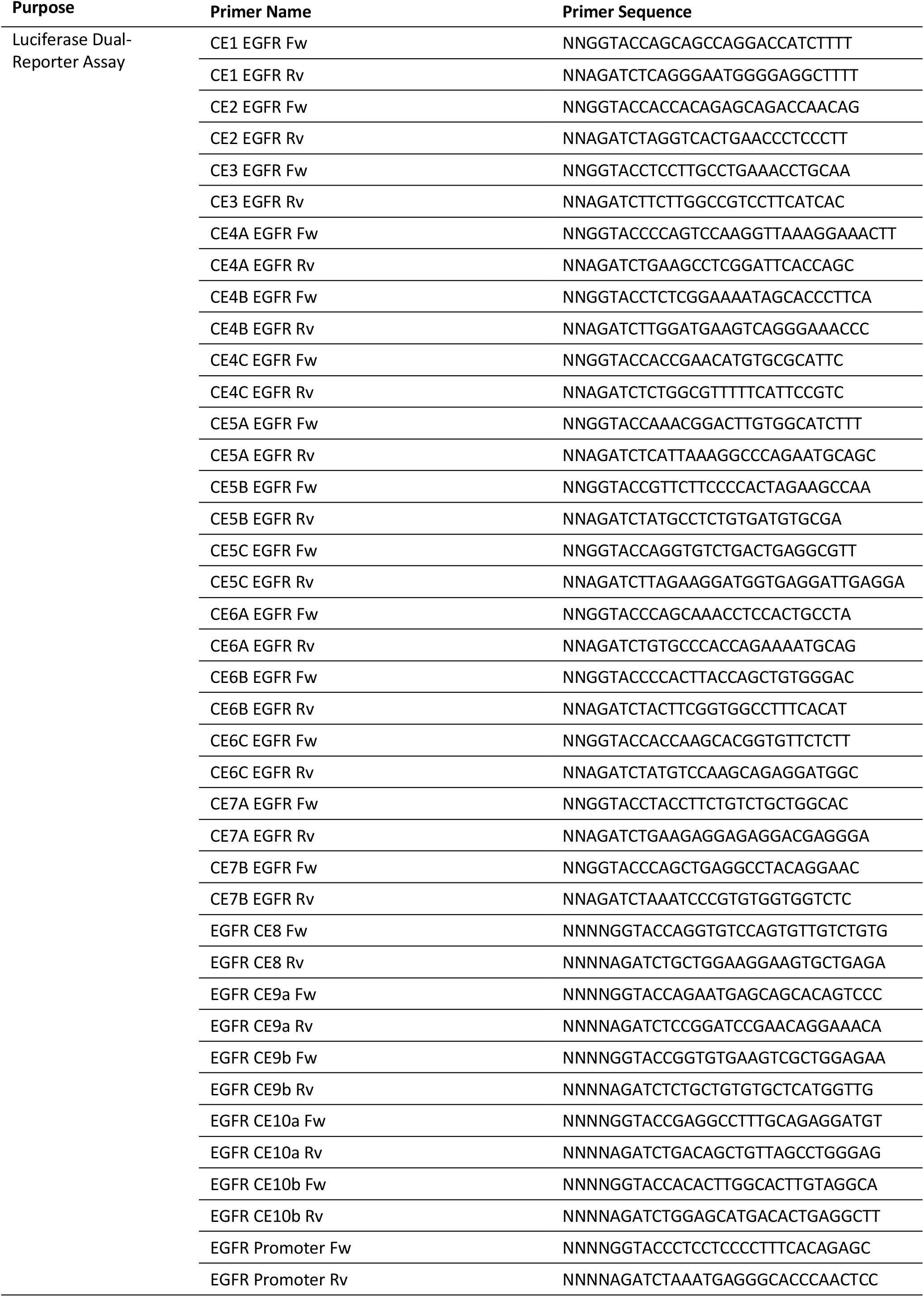

**Supplementary Table S2.**
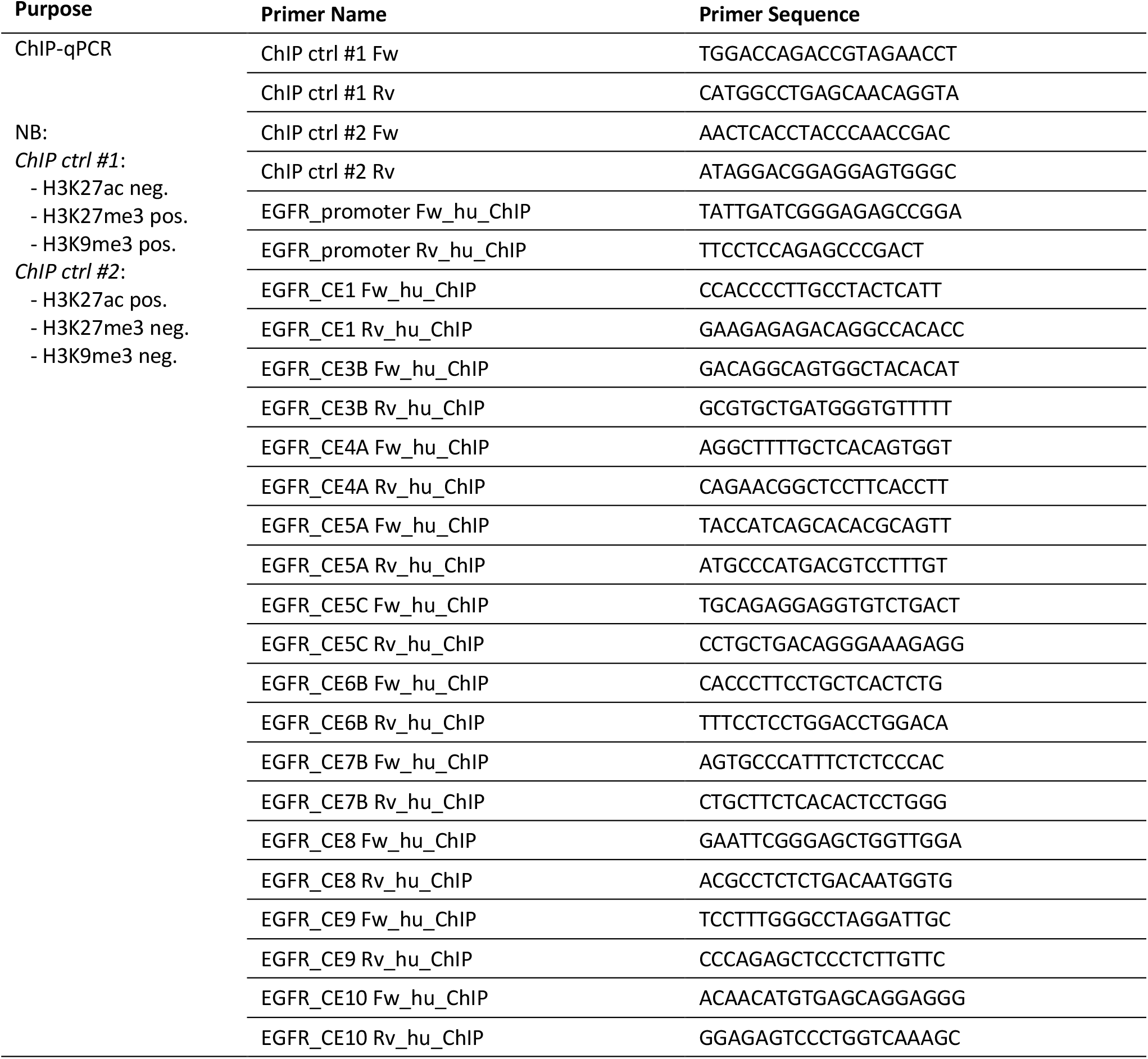

**Supplementary Table S3.**
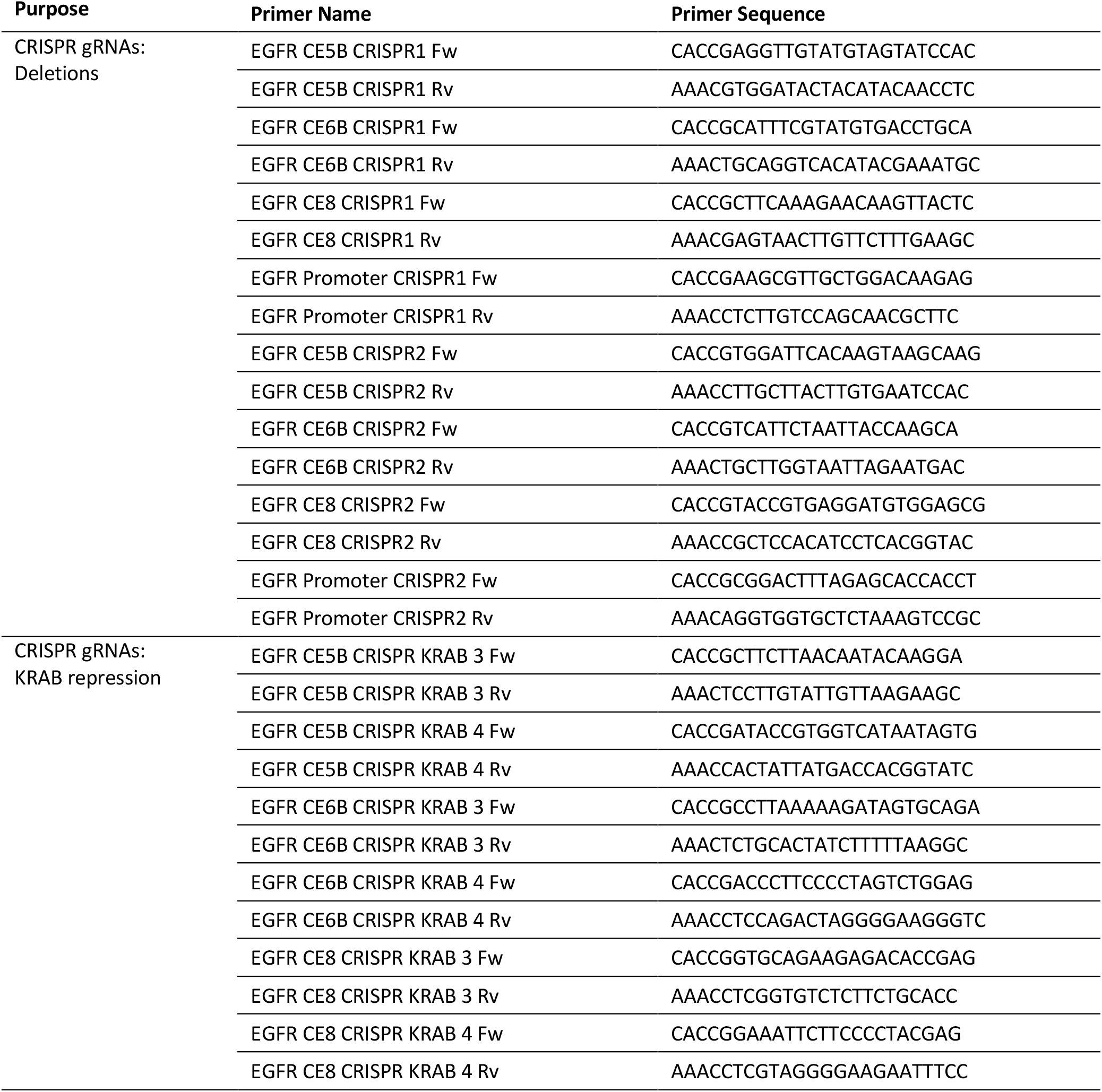

**Supplementary Table S4.**
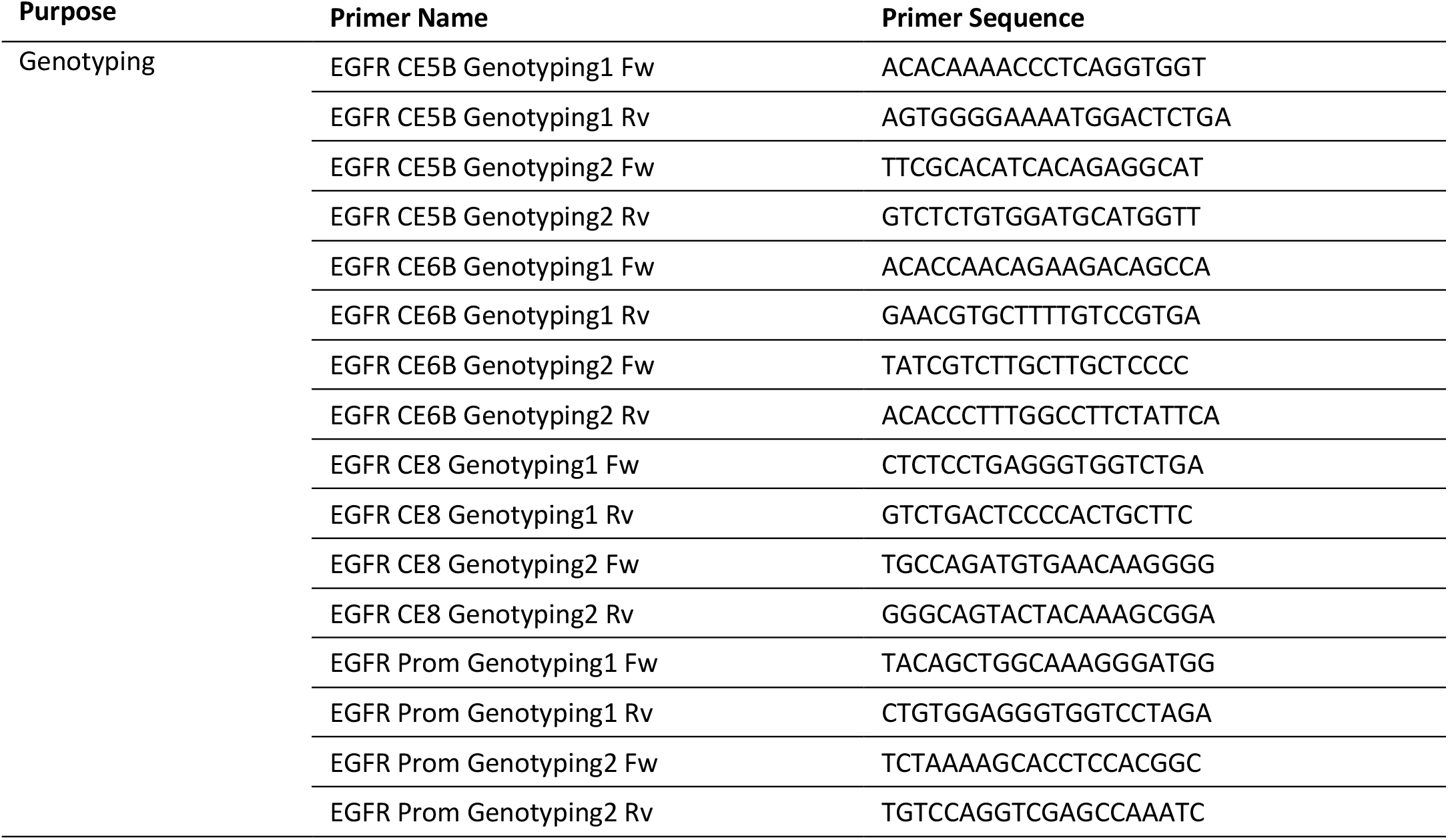

**Supplementary Table S5.**
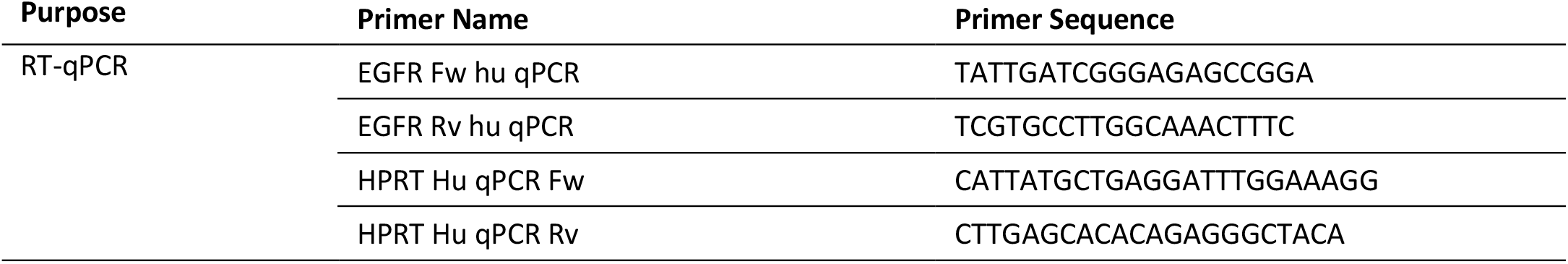

## Notes

CONFLICT OF INTEREST The authors declare no potential conflicts of interest.

### Competing Interest Statement

The authors have declared no competing interest.

